# Phylogenetic tree statistics: a systematic overview using the new R package ‘treestats’

**DOI:** 10.1101/2024.01.24.576848

**Authors:** Thijs Janzen, Rampal S. Etienne

## Abstract

Phylogenetic trees are believed to contain a wealth of information on diversification processes. Comparing phylogenetic trees is not straightforward due to their high dimensionality. Researchers have therefore defined a wide range of one-dimensional summary statistics. However, it remains unexplored to what extent these summary statistics cover the same underlying information and what summary statistics best explain observed variation across phylogenies. Furthermore, a large subset of available summary statistics focusses on measuring the topological features of a phylogenetic tree, but are often only explored at the extreme edge cases of the fully balanced or unbalanced tree and not for trees of intermediate balance.

Here, we introduce a new R package that provides speed optimized code to compute 54 summary statistics. We study correlations between summary statistics on empirical trees and on trees simulated using several diversification models. Furthermore, we introduce an algorithm to create intermediately balanced trees in a well-defined manner, in order to explore variation in summary statistics across a balance gradient.

We find that almost all summary statistics are correlated with tree size, and it is difficult if not impossible to correct for tree size, unless the tree generating model is known. Furthermore, we find that across empirical and simulated trees, at least two large clusters of correlated summary statistics can be found, where statistics group together based on information used (topology or branching times). However, the finer grained correlation structure appears to depend strongly on either the taxonomic group studied (in empirical studies) or the diversification model (in simulation studies). Nevertheless, we can identify multiple groups of summary statistics that are strongly and consistently correlated, indicating that these statistics measure the same underlying property of a tree. Lastly, we find that almost all topological summary statistics vary non-linearly and sometimes even non-monotonically with our intuitive balance gradient.

Therefore, in order to avoid introducing biases and missing underlying information, we advocate for selecting as many summary statistics as possible in phylogenetic analyses. With the introduction of the treestats package, which provides fast and reliable calculations, such an approach is now routinely possible.

## Introduction

Phylogenetic trees contain information on diversification in both the way nodes and tips are connected to each other (e.g. their topology) and in the sequence of branching events (e.g. the branching times). This source of information has been used extensively, for instance to infer underlying evolutionary processes such as speciation (Nee, 2001), extinction (Nee & Holmes, 1994), the interaction between traits and diversification (Herrera-Alsina et al., 2018), to reconstruct ancestral states (Pagel et al., 2004; Yu et al., 2015) and within the field of phylogenetic comparative methods (Revell & Harmon, 2022).

However, phylogenetic trees are hard to compare directly with each other due to their high dimensionality and comparison between trees is often made using lower dimensional summary statistics. Studies often focus on a subset of all available summary statistics, attempting to capture as much information as possible. This ranges from looking only at a single statistic (often a branch-times related statistic) ((Guimarães Fabreti & Höhna, 2023; Harmon et al., 2003; Liow et al., 2010), combining a topology-based and branch-times based statistic in an effort to cover as much information as possible with a very small selection of statistics (Hagen et al., 2015; Janzen & Etienne, 2017; Verboom et al., 2020), or using a large number of summary statistics in Approximate Bayesian Computation (Saulnier et al., 2017), Machine Learning (Ruffley et al., 2019) and Deep Learning (Lambert et al., 2023; Voznica et al., 2022). However, it remains unclear to what extent choices of used summary statistics are impacted by potential correlations and information overlap between statistics. An important first step towards an increased understanding has been taken by Tucker and colleagues, who investigated the choice of summary statistic for community ecology related studies, and identified several ‘anchor’ summary statistics, representing three important dimensions of a phylogenetic tree: richness, divergence and regularity (Tucker et al., 2017). For richness, they selected phylogenetic diversity, for divergence, mean pairwise distance and for regularity, variance of pairwise distance. However, their approach was geared towards community assembly whereas we focus on clade diversification.

A subset of all summary statistics focuses on measuring the balance of a tree (see for a comprehensive review (Fischer et al., 2021). Balance of a tree reflects the distribution of daughter tips in different parts of the phylogenetic tree, mimicking the physical act of balancing a tree. Generally, there is consensus on what constitutes a fully balanced tree: where each branch bifurcates into two daughter branches until reaching the present time. Analogously, a fully imbalanced tree reflects a tree where all daughter branches stem from the same parental lineage, such that all branches do not have their own daughter branches, generating a so-called ‘caterpillar’, ‘ladder’-like or ‘comb’ tree (Figure 1). Balance statistics then, maximize their value on a fully balanced tree, and minimize their value on a fully imbalanced tree. Similarly, imbalance statistics do the inverse. What remains unexplored however, is how summary statistics capture intermediately balanced trees, e.g. can we use summary statistics to identify how close we are to a fully (im)balanced tree?

**Figure 1.**
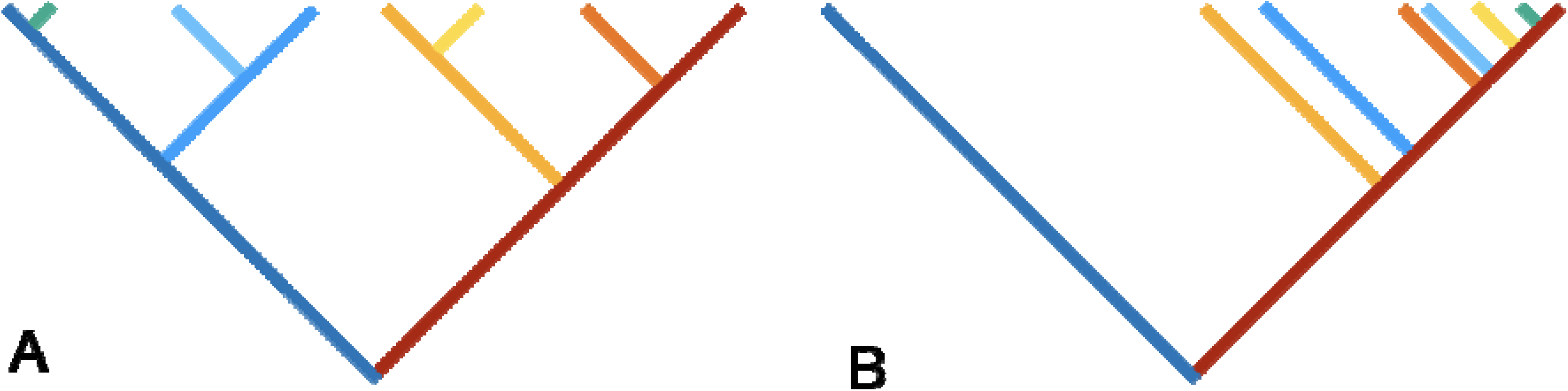
A fully balanced (A) or fully imbalanced (B) tree of 8 tips, highlighting the two crown lineages (blue and red), and their respective daughter lineages, where daughter lineages are colored depending on their parent lineage (e.g. all daughter lineages stemming from the blue crown lineage are colored in a blue-spectrum color). Both trees have the same sequence of branching times and same branch lengths, and tree B has been constructed from tree A by re-attaching all branches to the dark red crown lineage.

Furthermore, it remains unclear to what extent different balance metrics measure the same aspects of balance of a phylogenetic tree.

Here, we introduce the R package treestats, that computes 53 existing summary statistics for phylogenetic trees, allowing for a quick and user-friendly analysis of the interrelationship between summary statistics. We explore and investigate correlations between summary statistics on both empirical and simulated phylogenetic trees. Furthermore, the package computes a new balance statistic (the imbalance steps statistic) we introduce here, alongside algorithms to generate trees of intermediate balance, and we study how balance statistics vary with varying degrees of intermediate balance.

## Methods

We introduce the R package ‘treestats’, which computes 54 summary statistics for phylogenetic trees. For each statistic, we have developed a fast (Figure S2), C++ based, implementation from scratch, that generates a speed-up of often several orders of magnitude (Figure S3).

### Summary Statistics

We describe each summary statistic, where we classify them using the general source or type of information used per statistic.

#### Node statistics

Firstly, we collected a number of statistics summarizing properties of nodes, in strictly binary, bifurcating, trees. Each node indicates the branching event into two daughter lineages. Here, we use the notation ‘L’ and ‘R’ to indicate the number of tips attached to the ‘Left’ and ‘Right’ daughter branch. Summing over all nodes in the tree, we have the following statistics, indicating the calculation per node:

1. Colless index, abs (L – R) (Colless, 1982)
2. Blum statistic, log(L+R-1) (Blum & François, 2006)
3. Rogers J index, L != R (Rogers, 1996)
4. equal weights Colless index, abs(L – R) / (L + R – 2) (Mooers & S. B. Heard, 1997)
5. Number of pitchforks, L + R = 3 (Kendall et al., 2018)
6. Number of cherries, L + R = 2 (McKenzie et al., 1999)
7. Stairs measure, the fraction of nodes with (L != R) (Norström et al., 2012)
8. Stairs2 measure, the average of min(L, R) / max(L, R) (Norström et al., 2012),
9. ILnumber, nodes with a single tip child, e.g. either L = 1 or R = 1 (Kendall et al., 2018)
10. Rquartet index, (Coronado et al., 2019)
11. I statistic, (max(L, R) – (L + R) / 2) / (L + R – 1 - (L + R) / 2) (Fusco & Cronk, 1995; Purvis et al., 2002)
12. J^1^, -(L/(L+R)) log(L/(L+R)) (Lemant et al., 2022)
13. Beta (beta_dist(L, R)) (Aldous, 1996)
14. Average ladder measure, fraction of nodes with either R = 1 or L = 1 and that are connected with each other (e.g. forming a ladder) (Kendall et al., 2018)
15. Maximum ladder statistic, longest chain of connected nodes with either R = 1 or L = 1 (e.g. forming the longest ladder) (Kendall et al., 2018)

#### Depth statistics

The depth of a node (or tip) is defined as the distance between the root (or crown) and that node (or tip). Statistics using the depth are:

16. Maximum depth, measuring the maximum path (distance) between the tips and the root (Colijn & Gardy, 2014).
17. Sackin index, the sum of depths across all tips (Sackin, 1972)
18. Average leaf depth, the average depth of all tips (Shao & Sokal, 1990)
19. Variance of leaves’ depth, the variance in tip depths (Coronado et al., 2020).
20. B1, the sum of the inverse of all node depths (Shao & Sokal, 1990)
21. B2, the sum of the depth of each node divided by 2^depth^ (Shao & Sokal, 1990).
22. Symmetry Nodes index, the number of internal nodes that are not symmetry nodes, e.g. nodes from which the two subtrees have the same tree shape. Tree shape is measured as the distribution of node depths across all descendent nodes (Kersting & Fischer, 2021).

The width of a tree represents the number of occurrences of each depth in a tree. Summary statistics making use of this are:

23. Maximum width, the most common depth (Colijn & Gardy, 2014).
24. Maximum difference in widths, the maximum of the differences in occurrences between consecutive widths, where the widths are ordered by size (Colijn & Gardy, 2014).

#### Distance matrix

The distance matrix represents all pairwise distances between tips (and potentially nodes), either expressed in the sum of branch lengths, or in the number of edges required to traverse the shortest path between the pair. Summary statistics based on this distance matrix are:

25. Total cophenetic index, the sum of the distances to the lowest common ancestor between two tips, expressed in number of edges (Mir et al., 2013).
26. Area per pair index, the sum of the number of edges on the path between all pairs of tips (Lima et al., 2020).
27. Mean pairwise distance (mpd), the mean distance between all pairs of tips in units of branch length (Tsirogiannis et al., 2012; Webb et al., 2002)
28. Variance in pairwise distance statistic, the variance in distance between all pairs of tips in units of branch length (Webb et al., 2002).
29. J statistic, the intensive quadratic entropy given by the average distance between two randomly chosen species, calculated as the sum of all pairwise distances divided by S^2^, where S is the number of tips of the tree (Izsák & Papp, 2000)
30. Mean nearest taxon distance, the mean distance to the nearest tip (Webb et al., 2002).
31. Phylogenetic species variability index (psv), the sum of all pairwise relationships d_ij_, where d_ij_ = 0.5 * (d_ii_ + d_jj_ + d_ij_), where d_xx_ indicates the distance of tip x to the root (Helmus et al., 2007).

The distance matrix also serves as a starting point for measuring the Laplacian spectrum. The starting point here is a Laplacian matrix, which is the distance matrix of the phylogenetic tree, but on the diagonal the zero distances are replaced by the negative row sums. Using the Laplacian matrix, the Laplacian spectrum statistic then calculates the eigenvalues and eigenvectors of the Laplacian matrix (Lewitus & Morlon, 2016). We identify four properties of interest here:

32. Principal eigenvalue of the Laplacian matrix (laplace_spectrum_e in all figures)
33. Eigen gap of the Laplacian matrix, e.g., the position of the largest difference between eigen values, indicating the number of modalities in the tree (laplace_spectrum_g in all figures)
34. Asymmetry of the distribution of eigenvalues of the Laplacian matrix (laplace_spectrum_a in all figures)
35. Peakedness of the eigenvalues distribution of the Laplacian matrix (laplace_spectrum_p in all figures)

#### Network science

Summary statistics that take inspiration from network science, consider phylogenetic trees in their unrooted form, and use network properties to calculate properties of the trees. These are:

36. Wiener index, the sum of the shortest path lengths across all tips of the tree (Chindelevitch et al., 2021).
37. Diameter the maximum of all shortest paths between all tips of the tree (Chindelevitch et al., 2021). When taking branch lengths into account, this is equal to twice the crown age in an extant tree (see below).
38. Maximum betweenness, the maximum value of betweenness for each node *u*, where betweenness is measured as the shortest path between nodes *v* and *w*, going through focal node *u* (Chindelevitch et al., 2021).
39. Maximum closeness, the inverse of farness, where farness is the sum of distances from a node to all the other nodes in the tree (Chindelevitch et al., 2021).
40. Eigenvector, the principle component of the Perron-Frobenius eigenvector of the adjacency matrix of the tree (Chindelevitch et al., 2021).

#### Branching times

The sequence of branching times mainly holds information about the timing of branching events, and disregards any information held in the topology. Summary statistics based branching times are:

41. Crown age, the maximum branching time
42. Tree height, the maximum branching time plus the root branch length (aka stem age).
43. Gamma, the relative position of internal nodes, with respect to the root and the tips (Pybus & Harvey, 2000)
44. Pigot’s rho, the slope of diversification in the first and second half of the tree, corrected for tree size (Pigot et al., 2010).
45. nLTT, the difference in sequence of branching times between two trees; this measures the alignment of diversification between two trees (Janzen et al., 2015).

For independent comparison, the treestats package introduces the function ‘nLTT_base’, which compares a focal tree with a non-diversifying tree with only two lineages.

#### Branch lengths

Related to branching times, a number of summary statistics derive from the distribution of branch lengths of a tree, ignoring the associated topology:

46. Phylogenetic diversity, the sum of branch lengths (Faith, 1992; Richter et al., 2021)
47. Mean branch length, the sum of branch lengths divided by the number of branches (Janzen & Etienne, 2017)
48. Variance of branch lengths (Saulnier et al., 2017)

Saulnier et al further subdivide branch lengths into two groups: external branch lengths, e.g. lengths of branches attached to the tips of the tree, and internal branch lengths, e.g. lengths of branches only attached to internal nodes of the tree. For these two groups, they then introduce the average and variation as statistics, to end up with:

49. Mean branch length of external branches (Saulnier et al., 2017)
50. Mean branch length of internal branches (Saulnier et al., 2017)
51. Variance of branch length of external branches (Saulnier et al., 2017)
52. Variance of branch length of internal branches (Saulnier et al., 2017)

As a last statistic, we have:

53. Number of tips

We further introduce a new statistic, explained in the next section:

54. Imbalance steps

## A new statistic: Imbalance steps

For every reconstructed phylogenetic tree, we can represent it such that the tree, starting at the crown of the tree, consists of two main branches (crown lineages A and B), from which daughter branches branch off, which in turn might generate their own daughter branches. This looks very much like a classic cladogram.

At one extreme we have the fully balanced tree: assuming that a tree has 2^n^ = N lineages, in each half of the tree we then find N/2 tips, of which log_2_(N / 2) are direct daughters of the crown lineages, and N/2 – log_2_(N/2) are daughter lineages of daughter lineages etc. This yields a fully balanced topology, with the two crown lineages functioning as main ‘stems’ of the tree (figure 1A). At the other extreme we have the fully imbalanced tree, where again we have the two main branches, but now find that all daughter branches are attached to one of the crown lineages, and that all daughter branches are terminal branches, i.e., they do not have daughter lineages themselves (figure 1B).

If we keep track of how branches are attached to each other, we can transform the fully balanced tree into the fully imbalanced tree in a stepwise fashion, by re-attaching branches to one of the two crown lineages, until all branches are terminal branches and attached to the same crown lineage.

Then, we define the imbalance steps statistic (statistic 54) as the number of steps it takes to transform a tree into a fully imbalanced tree, where high values indicate a very balanced tree, and low values indicate a very imbalanced tree.

For a fully balanced tree of N tips, the number of steps required to transform the tree into a fully imbalanced tree is N – (log_2_(N / 2) + 2), where log_2_(N / 2) is the number of daughter lineages already connected to the crown lineages accepting re-attached branches, and 2 is the two crown lineages. This simplifies to: N – log_2_(N) – 1, and we can use this expression to normalize the observed number of required steps by tree size.

Although the total number of steps required is straightforward to calculate, observing and studying trees of intermediate balance, i.e., that are not fully (im)balanced is less straightforward: there are many different move orders possible to go from a fully balanced to a fully imbalanced tree. Here we explore a number of move algorithms, that can be categorized firstly into two groups, based on the set of potential branches that the algorithm chooses from: 1) group A that re-attaches any branch, and 2) group T that re-attaches only terminal branches (e.g., branches that have no daughter branches). Then, within each group, there are three ways to determine which branch from all available branches to pick: R) a branch is selected randomly, Y) the youngest (shortest) branch is selected or O) the oldest (longest) branch is selected. Combining these two we obtain six selection algorithms (A-R, A-Y, A-O, T-R, T-Y & T-O). However, both TY and AY pick from the same branch set (when selecting the youngest branch, this is always a terminal branch), and thus we obtain five unique selection algorithms: A-R, A-O, T-R, T-Y and T-O.

Please note that these algorithms determine the order in which branches are moved, but do not influence the total number of steps required (e.g. the imbalance steps statistic).

### Empirical data

To assess correlations across statistics, we assess all summary statistics on a dataset of 215 phylogenetic trees collected on a family level (Condamine et al., 2019). These trees vary in size from 10 to 680, and are broadly grouped into four taxonomic groups: birds (129 trees), mammals (66 trees), squamates (11 trees) and amphibians (9 trees). We computed for each tree all 54 summary statistics.

To correct for phylogenetic relatedness between families and/or taxonomic groups, we constructed a ‘supertree’ spanning all families using the R package datelife (O’Meara et al., 2023). We then calculated correlations between all statistics correcting for tree size and phylogenetic correlation.

To obtain PCA estimates we performed phylogenetic corrected PCA using the R package phytools (Revell, 2012)

### Simulated data

We explored several aspects of phylogenetic summary statistics by applying them to simulated trees. We simulated trees using a number of different diversification models. Unless stated otherwise, all trees were simulated conditional on having 300 tips and a crown age of 10 My. The diversification rate was set to 0.5, with either no extinction and speciation at 0.5, or with extinction set to 0.1 and speciation set to 0.5 + 0.1 = 0.6, thus keeping the rate of diversification the same as without extinction.

We explored the following models:

- Constant Rates Birth-Death model
- Diversity-Dependent Diversification model (Etienne et al., 2012), with K set to 300 + 20% = 360.
- Protracted birth-death model (Etienne & Rosindell, 2012) with λ (speciation completion rate) at 10, no incipient speciation and the extinction rate of good and incipient species equal.
- Binary State dependent diversification model (BiSSE) (FitzJohn, 2012) with b_0_ and μ_0_ equal to the abovementioned speciation and extinction rate, and b_1_ and c_1_ equal to 1.5 times b_0_ and μ_0_. We set the transition rates between states, q_01_ and q_10_, both equal to 0.1.

Parameter values for each diversification model were chosen in order to emphasize (but not exaggerate to the extreme) the different peculiarities of each model.

#### Tree Size

To study the effect of tree size and explore the effectiveness of tree size normalization across statistics, we simulated trees with a varying number of tips across the four models, i.e. the number of tips N varied in 10^[1, 1.1, 1.2, …, 3]. We kept the speciation rate constant and varied the crown age in order to allow for the diversification process to be able to reach the target number of tips, where the crown age T was given by: 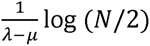.

#### Correlations

For each of the models, with or without extinction, we simulated 100.000 trees and calculated the corresponding summary statistics. Using these statistics, we calculated the Pearson rank correlation (R) of the summary statistics. We then constructed a distance matrix across summary statistics based on 1 – abs(R). Using this distance matrix, we clustered summary statistics together using an unweighted pair group method with arithmetic mean (UPGMA) algorithm.

We performed this analysis for each diversification model separately, but also for the entire dataset, e.g. for all 1.200.000 trees combined (focusing on only trees with extinction).

#### Consistent correlations

To identify combinations of summary statistics that consistently correlate with each other, we measured the mean absolute correlation and the standard deviation across the absolute correlation across the four taxonomic groups (Mammals, Birds, Amphibians, Squamates) and the four diversification models (BD, DDD, PBD, SSE). Then, we selected the statististics with very low standard deviations (< 0.05) and high correlations (> 0.9).

## Results

### Empirical data

We find that across the empirical datasets, PCA analysis does not separate the different taxonomic groups (i.e. amphibians, birds, mammals and squamates) based on summary statistics of the used phylogenetic trees, supporting further analysis for all trees together (Figure 2C). Furthermore, we find that the first five axes of the PCA analysis explain 80.4% of the observed variation (Figure 2B), suggesting that careful selection of about five summary statistics might capture the large majority of observed variation across empirical trees. When we break down the contribution of each summary statistic for the first five PCA axes, we find that the first PCA axis (45.9% variation explained) and the second PCA axis (13.1% variation explained), are mainly driven by topology-based summary statistics such as Rogers, B1, Stairs and equal weights Colless. The third (8.1 % variation explained) and fourth PCA axis (7.5% variation explained) are mainly driven by branch length-based summary statistics including crown age, pigot’s rho, Mean Pair distance and variation in branch length. The fifth PCA axis (5.9 % variation explained) Is mainly driven by topology associated statistics again, including maximum closeness, B2 and rquartet.

**Figure 2.**
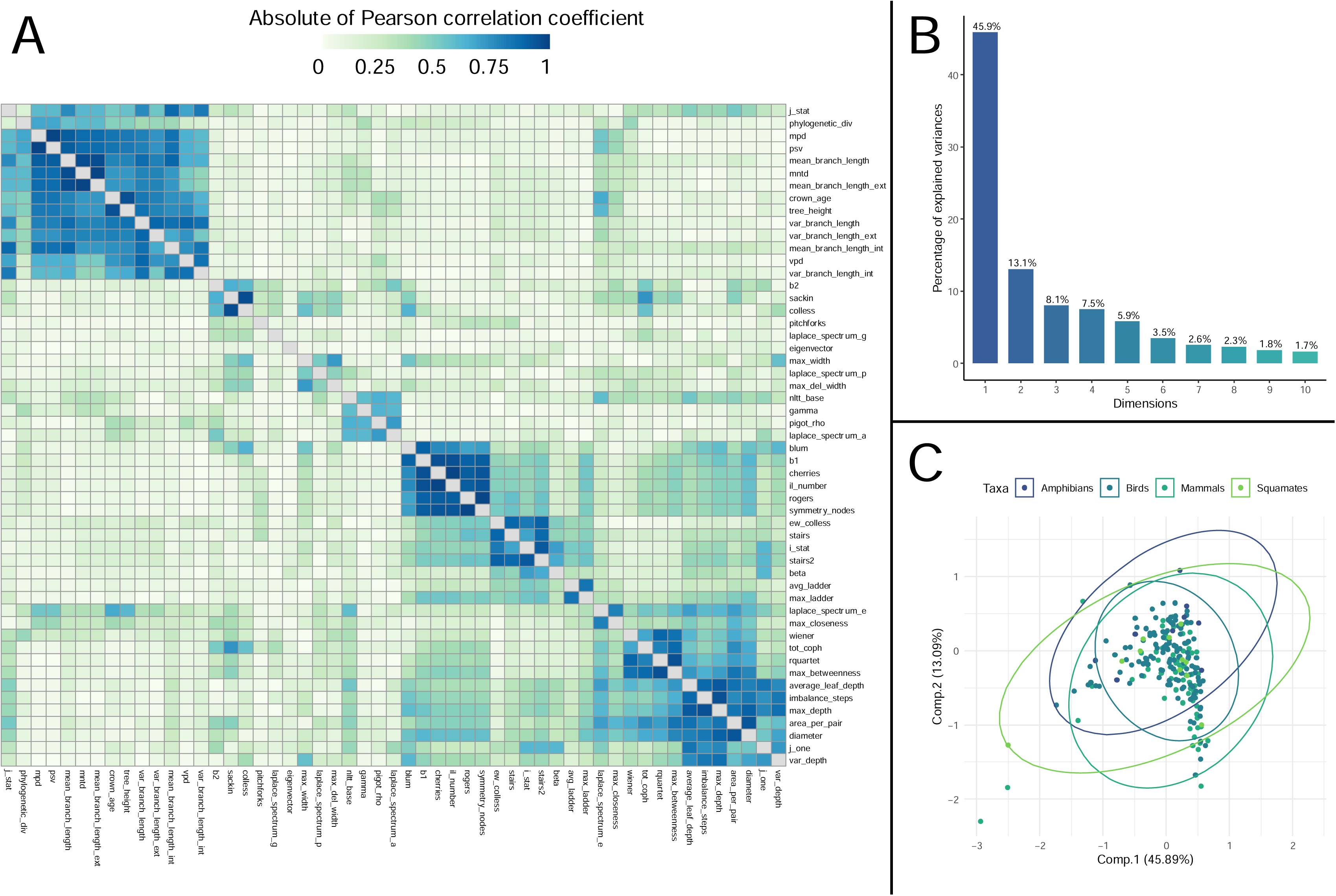
**A**: Absolute of Pearson Rank Correlation between summary statistics calculated for 215 phylogenetic trees sampled across the tree of life. Correlations are corrected for phylogenetic relatedness and tree size. **B:** percentage of explained variance of a PCA analysis of all 54 statistics on the empirical phylogenies, showing that the first five dimensions explain at least 80 % of the observed variance. **C:** First two components of this PCA analysis, where dots are colored according to the respective taxonomic group studied, showing that across the tree of life, phylogenetic tree properties, although variable, tend to share many characteristics.

Using the obtained correlations, we constructed a correlation dendrogram, which groups statistics that share high similarity (i.e. that are strongly correlated, either positively or negatively) together (Figure 3). We then grouped together statistics into larger clusters when statistics within a group at least had an absolute correlation of 0.2. For all taxonomic groups together, we find (Figure 3) one small cluster (gamma, nLTT, pigot’s rho and Laplacian spectrum Asymmetry) that combines statistics related to branching times. Grouping next to this cluster, we recover a larger cluster including crown age, mean pair distance and variation in branch length that only includes statistics based on branch lengths. Next to this, we recover two larger clusters with statistics based on topology: one cluster including the Sackin and Colless index, and one cluster including the stairs, cherries, J^1^ and Total cophenetic index. Lastly, we find that pitchforks and eigenvector do not group strongly within either of these three larger clusters. The other panels in figure 3 show clustering of statistics within each taxonomic group. These show that for mammals, the clusters recovered in the overall dataset are well retained and still cluster together. For birds we find that the topology-based cluster associated with the Colless index is broken up. For Amphibians and Squamates we find that none of the clusters retain their clustering.

**Figure 3.**
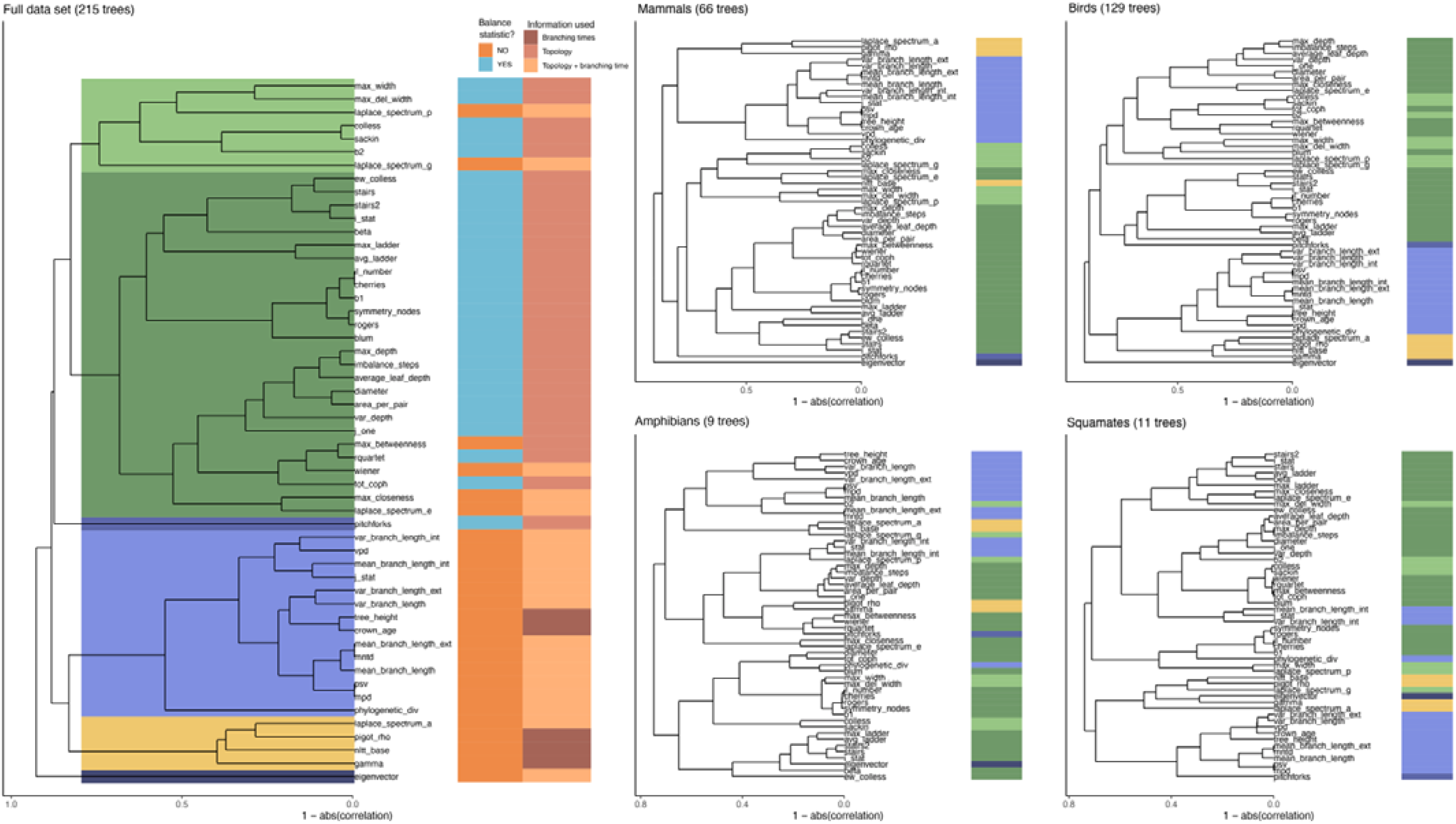
Correlation dendrograms shown for the entire empirical dataset (left) and for each taxonomic group separately. Correlation dendrograms are based on 1 – abs(correlation). All correlations are corrected for size and phylogenetic signal. Colors spanning the dendrogram show clusters based on a cut-off of a minimum absolute correlation of 0.2. These spanning colors return next to the tips in each of the taxonomic group-based correlation dendrograms, to indicate how well the overall clustering is preserved within each taxonomic group. Colors next to the tips in the entire dataset dendrogram indicate whether the statistic in question is defined as a balance statistic (yes = blue, no = orange), and the type of phylogenetic information required to calculate the relative statistic (only topology, only branching times, or both topology and branching times).

### Simulated data: correlations

To avoid effects of tree size, we simulated trees under different diversification models, keeping tree size (and crown age) constant. Then, we constructed correlation matrices for the different models (Figure 5). Similarly, to the empirical data, we recover three large clusters of statistics, but this time with only one singleton statistic that does not belong in any of these large clusters: max_betweenness. Again, we find that all branching time related statistics cluster together. However, in contrast to the empirical data, we find that the topology-related statistics can be subdivided into two groups, one associated with the Sackin and Colless indices, and one associated with the cherries, pitchforks and stairs statistics. Across the four diversification models, we find that the topology cluster including cherries and stairs is largely retained, although with the PBD model, two statistics appear within the cluster. Similarly, the cluster including the Colless and Sackin indices is also usually largely retained, although often one or several statistics cluster differently, depending on the diversification model. Lastly, the branching times cluster is often subdivided into two subsections: one including statistics correlating with mean_branch_length and one including statistics correlating with the mean pair distance.

### Simulated data: tree size

Empirical trees are an invaluable source of information, but are all different in size. This tree size variation may cause spurious correlations (Figure 4). For some statistics, normalization terms are available that correct for this, but we suspected that these may be insufficient. To test this, we simulated trees of different sizes under a range of different diversification models. We find (Figure 4) that only the cherries and pitchforks statistics are able to correct for tree size, regardless of the underlying model. The area per pair, Colless, Sackin and total cophenetic index can also correct for tree size, but only if the model used to simulate the tree matches that for the expectation, which is a Yule model. In all other cases, we find that tree size, even after correction, still correlates with the summary statistic.

We recover a similar pattern when we measure summary statistics through time in a tree (Figure S4), where we observe that the cherries and pitchforks statistics show little variation across the age of the tree, whereas other statistics change considerably – although those changes are of course also due to diversification dynamics changing over time, depending on the diversification model chosen.

**Figure 4.**
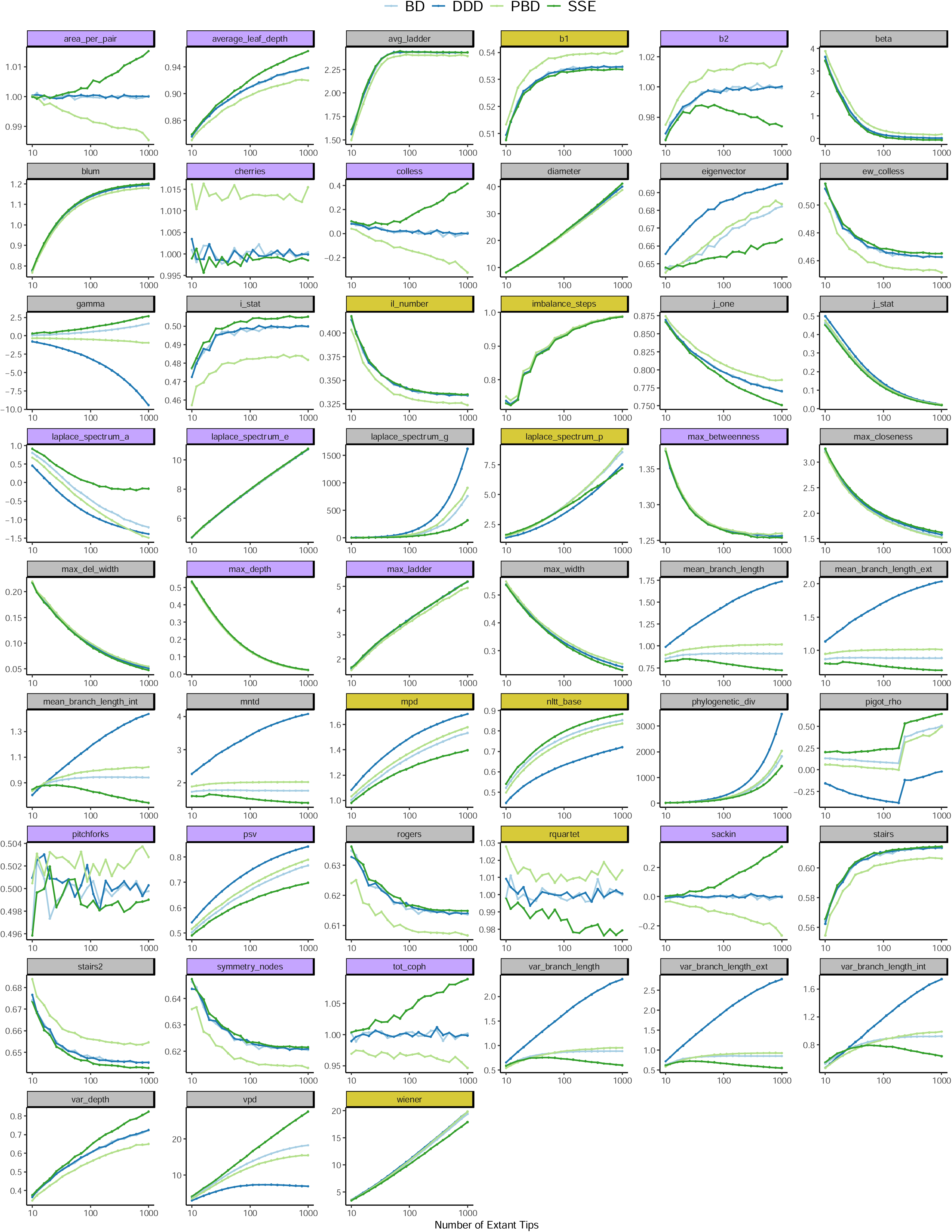
Observed variation of tree statistics across differently sized trees. Gold colored statistics use the number of tips to normalize the summary statistic, Purple colored statistics use the expectation under the Yule model to normalize the summary statistic. Grey colored tree statistics do not provide the option to normalize for tree size. Shown are the average tree statistic values across 10.000 replicates per model, per tree size, for four diversification models.

### Consistent correlations

The larger scale correlation structure appears quite variable, but within these larger clusters, we identify multiple statistics that are very strongly and consistently correlated with each other, indicating a strong overlap in information content. Table 1 shows that the Colless and Sackin indices tend to correlate strongly with each other (mean correlation of 0.96), even more so when only considering simulated trees of identical size (mean correlation of 0.98). Interestingly, we find that in simulations, there is a larger tight cluster around the Sackin and Colless indices, including the J^1^ index, Average leaf depth and the Total Cophenetic index. It not clear why these statistics correlate so strongly, and why only on simulated trees. This is in contrast to the strong correlation of mean nearest taxon distance (mntd) and mean branch length of external branches, where the distance to the mean nearest taxon is mathematically equivalent to twice the length of the attached external branch, explaining the perfect correlation. Similarly, the mpd and psv statistics have previously been shown to be equivalent (Tucker et al., 2017), but we unexpectedly found that the Wiener statistic (not stemming from community assembly theory, but from network theory!), and the Entropy J statistic also strongly correlated with these two statistics.

**Table 1.** Identified tight clusters of strongly and consistently correlated summary statistics. Top part: Tight clusters across all observed tree collections (e.g. the four taxonomic groups, and the four diversification models), bottom part: tight clusters only observed in simulated tree collection. Every row indicates a separate tight cluster identified across all datasets, starting with a first statistic and then reporting each strongly correlating other statistic, where numbers between brackets represent the mean absolute correlation and standard deviation.

| *Across all trees* |  |  |  |  |
| --- | --- | --- | --- | --- |
| First Statistic | Second statistic | Third statistic | Fourth Statistic | Fifth Statistic |
| cherries | IL_number (1, 0.0) | b1 (0.93, 0.03) |  |  |
| Colless | Sackin (0.96, 0.05) |  |  |  |
| imbalance_steps | max_depth (0.99, 0.02) |  |  |  |
| Rogers | symmetry nodes (0.98, 0.009) |  |  |  |
| mntd | mean_branch_length_ext (1, 0) |  |  |  |
| mpd | psv (1, 0) |  |  |  |
| I_stat | stairs2 (0.94, 0.02) |  |  |  |
| **Simulations only** |  |  |  |  |
| Colless | Sackin (0.98, 0.002) | j_one (0.98, 0.001) | average_leaf_depth (0.98, 0.002) | tot_coph (0.9, 0.005) |
| Rogers | symmetry nodes (0.97, 0.0006) | stairs (1, 0) | stairs2 (0.92, 0.001) |  |
| cherries | ilnumber (1, 0) | b1 (0.92, 0.001) | ew_colless (0.91, 0.0008) |  |
| mpd | psv (1, 0) | j_stat (1, 0) | wiener (0.97, 0.05) |  |
| nLTT | phylogenetic diversity (0.999, 0.003) | mean_branch_length (0.999, 0.003) |  |  |
| Beta | Blum (0.91, 0.01) |  |  |  |

### General information content

When we look at how each summary statistic correlates with all others, we find that there is quite a difference between statistics, with some statistics having relatively high average correlation values (Figure 6), whereas others appear to hardly correlate with any of the other statistics, indicating they contain information not captured by other statistics. For empirical trees, we find that the eigenvector, mpd and gamma statistics perform especially well in this respect, and also three out of four components of the Laplacian spectrum associated summary statistics show low median correlation values. For simulated trees, again the eigenvector and gamma statistic show low average correlation values, but so do the ladder statistics (avg and max) and pigot’s rho.

Interestingly, those statistics that group outside of the main clusters in the previously mentioned correlation dendrograms (Figures 3 and 5), do not necessarily have the lowest average correlation. Clustering in the dendrograms is determined using an UPGMA algorithm, which first groups statistics together based on their closest sister statistic. For max betweenness in the simulated trees, we for instance observe that although its median correlation is not the lowest of all summary statistics, it does not show any outliers, indicating that its maximum correlation is very low as well – hence it does not group very well with other statistics (as shown in Figure 5). This is in contrast to the gamma statistic, which, although having a low median correlation, does show a limited number of correlations that are much higher and which cause it to group within the branch length associated cluster.

**Figure 5.**
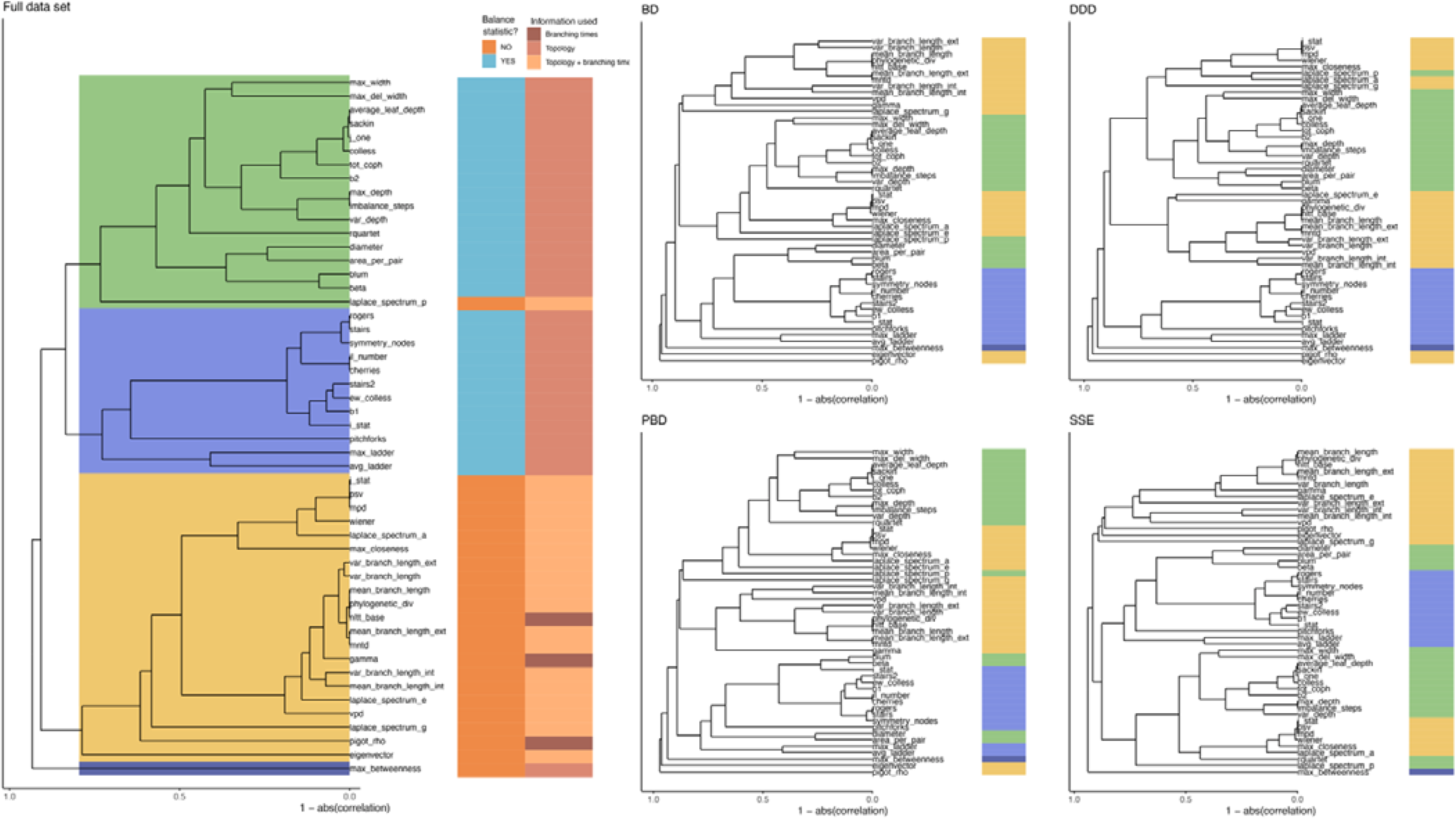
Summary statistics grouped together based on their correlations across 1,200,000 trees, consisting of 300,000 trees simulated per model (BD, DDD, PBD and SSE) with extinction. Colored rectangles indicate statistics grouped together based on a minimum absolute correlation of 0.2, please note that colors are arbitrary and do not correspond to colors in Figure 3. Colored rectangles at the right indicate in the first column whether the associated statistic is an (im)balance statistic or not and in the second column the type of information required to calculate the statistic.

**Figure 6.**
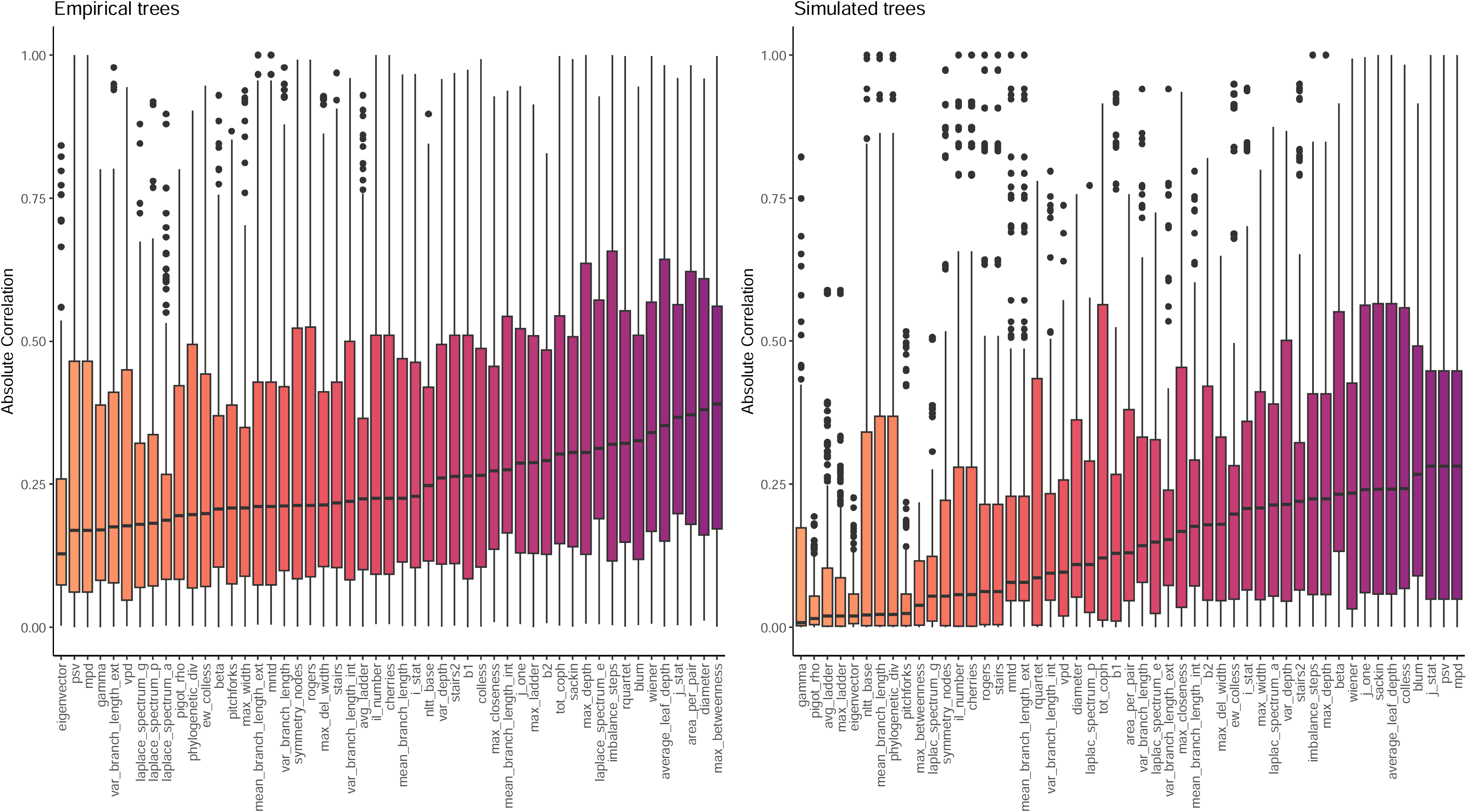
Absolute correlation coefficients with all other summary statistics, per summary statistic, for empirical trees (left panel, same trees as in figure 2) and simulated trees (right panel, same trees as in figure 4). Statistics are sorted by their median absolute correlation value. High values indicate that a statistic has high overlap with other statistics, whereas low values indicate that a statistic contains a high amount of unique information about the phylogenetic tree, not captured by other statistics.

### Intermediate balance

Using the algorithms described to create intermediately imbalanced trees, we explored (im)balance statistics on intermediately balanced trees. We find that almost all statistics vary non-linearly between the two known extreme values (Figure 7) (e.g. the perfectly balanced and completely imbalanced tree). Only the diameter and max depth statistic show a linear relationship with intermediate balance. Furthermore, we find strong variation across the different imbalancing algorithms, suggesting that capturing intermediate balance is a non-trivial endeavor and that intermediate balance may, depending on the tree topology, lead to very different summary statistic values. For the branch selection algorithm, we find that first selecting terminal branches as opposed to randomly selecting branches tends to lead to a slower approach of the value of the completely imbalanced tree. The specific type of branch (random, oldest, youngest) seems to matter a great deal as well, where selecting older branches causes larger initial deviations.

**Figure 7.**
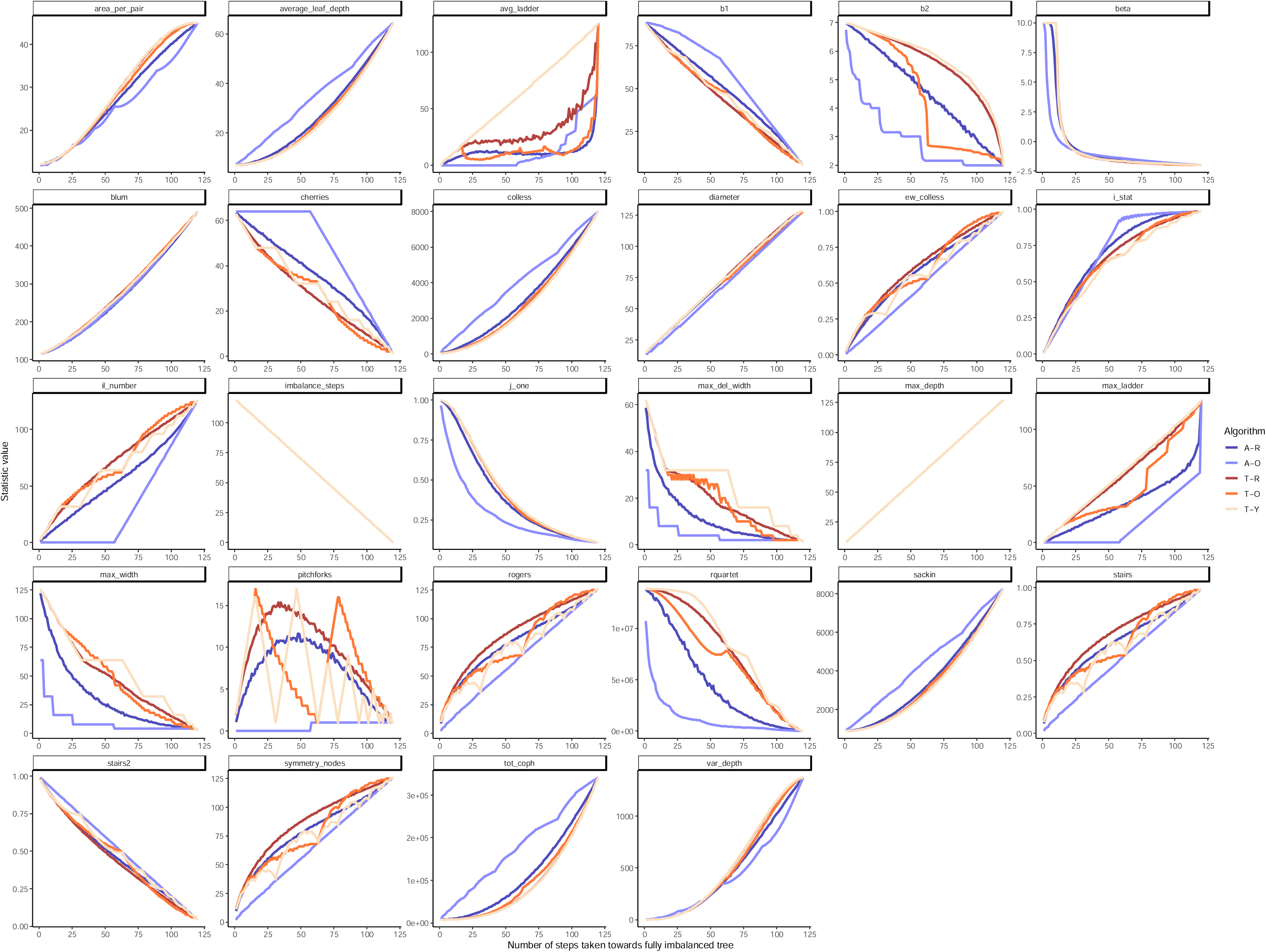
The average balance related summary statistic value of 100 replicate trees of 128 tips that are increasingly imbalanced using one of six algorithms available, using any branch, selecting at random (A-R), using any branch, selecting the youngest branch first (A-Y), using any branch, selecting the oldest branch first (A-O), using only terminal branches, selecting at random (T-R), using only terminal branches, selecting the youngest branch first (T-Y) and using only terminal branches, selecting the oldest branch first (T-O).

## Discussion

Here, we have introduced a new R package called ’treestats’, that provides fast (Figure S3, S4) and user-friendly code to quickly assess a wide range of phylogenetic summary statistics. We have used the package to analyze both empirical and simulated phylogenetic trees and unravel their interdependence. Firstly, we have found that almost all summary statistics are highly sensitive to effects of tree size, even after attempting to correct for tree size. Secondly, we have explored correlations across summary statistics on both empirical and phylogenetic trees, identified several larger clusters of closely related statistics and identified statistics with particularly low information overlap. Lastly, we have explored the variation of summary statistics with intermediately balanced trees, and found several summary statistics that vary non-monotonically with balance.

Many summary statistics correlate with tree size and therefore provide normalization functions to correct for this. However, we find that most of these corrections are inadequate and do not correct properly for tree size. In general, we find that there are two categories of normalizations available: corrections making use of the maximum value for a given tree size, and corrections using the average value expected under a diversification process. Corrections making use of the maximum value (such as for instance the Imbalance steps, b1 and Roger’s J statistic) fail to correct for the impact of the diversification process: for different tree sizes, the diversification process has a different expected summary statistic value, which is often different from the maximum obtainable value. Some other summary statistics recognize this (such as for instance the Sackin, b2 and Colless statistics) and provide normalization options depending on the Yule and PDA process. We find that when the generating process is close to these processes these normalizations work very well, but that when the generating processes are different, the normalization fails. Across all statistics, we find that only the cherries and pitchforks statistics are able to adequately correct for size, but for these statistics we do not find much variation across diversification models, suggesting low discriminatory power. Therefore, we recommend using tree size as a covariate when comparing summary statistics across trees of different sizes, instead of relying on self-normalization of a summary statistic.

Across empirical phylogenies, we recover four clusters of summary statistics that seem to cover the same information given their high correlation: 1) a cluster centered around balance related statistics such as Colless, Sackin and b2, 2) a large cluster centered around stairs and max_depth, 3) a cluster centered around branch length related statistics such as mean pair distance and variance of branch length and 4) a cluster centered around statistics related to branching times, such as gamma and nLTT. Furthermore, we recover two statistics that do not group within these clusters: pitchforks and eigenvector. The identification of these large clusters seems to suggest that the statistics within them tend to have increased overlap with each other, and that for comparisons across trees using only a limited number of statistics, statistics are best selected from different clusters, and/or those statistics that group outside these clusters. However, it should be noted that clustering within these clusters differed between taxonomic groups studied: within the group of birds, the cluster related to the Colless statistic became separated and grouped with other balance statistics. For the Amphibians and Squamates, we find that overall, the clustering is quite different from the clustering in Mammals and Birds, but we hypothesize that this could be confounded by the much lower number of trees analyzed in these taxonomic groups.

Across simulated trees, we recover three large clusters of summary statistics: 1) a large cluster centered around all branch length and branching time related statistics (such as Gamma, Mean Pair Distance and Mean Branch length), 2) a large cluster centered around balance statistics such as Colless, b2 and the Beta statistic and 3) a large cluster containing many pattern-based balance statistics, such as pitchforks, cherries and stairs statistics. For simulated trees, we only recover one outlier statistic, being the max betweenness statistic. Across diversification models we find that these clusters vary considerably, with the exception of cluster 3 that is consistently recovered across diversification models. Overall, we find a very clear grouping of balance related statistics on the one hand, and branch time or branch length related statistics on the other hand, indicating a fundamental divide between summary statistics. This pattern is much clearer for the simulated data than it is for the empirical data.

Comparing our results for our analysis on empirical data and on simulated data, we find that many patterns that arise for the empirical data, do not show up for the simulated data. This seems to suggest several things: firstly, the diversification models used in our simulated study are unlikely to be the generating processes for the empirical trees. Either because perhaps not all phylogenetic trees are the result of the same diversification process, or because the true diversification model was not amongst the models we explored. Secondly, in our simulation study we kept tree size constant, in order to avoid having to do a statistical correction for tree size in our results. However, the empirical data did have variation in tree size and we cannot exclude that perhaps the covariance with tree size (although corrected for) may have partially impacted the results. Lastly, the difference between empirical and simulation results indicates that information overlap between summary statistics is not independent of the tree diversification process, highlighting the importance of understanding the generating process of phylogenetic trees.

Before embarking on this study, we expected to find some statistics that were measuring (mathematically) the same properties of trees, irrespective of the process generating the tree. For some statistics, this was already known (such as the PSV and MPD statistic (Tucker et al., 2017)), but we expected to recover some more spurious relations as well. To our surprise, we recovered that for many statistics, such correlations were highly dependent on the generating model, indicating that those statistics did not have strong mathematical overlap. However, we also do identify a number of statistics that show very strong correlations with each other, regardless of taxonomic group or diversification model used, indicating high information overlap. The most interesting correlation group is a group consisting of the Sackin, Colless, J1, Total cophenetic and Average Leaf Depth statistics, which all correlate strongly with each other, indicating that they can be used interchangeably. For some other combinations of statistics, we only recover strong, consistent, correlations for simulated trees (e.g., imbalance steps and max depth). It is unclear why these correlations disappear in the empirical tree dataset, but these results again indicate that the diversification models used in our simulations are different from the processes that have generated the empirical trees.

Using the average correlation across either taxonomic groups or diversification models, we identify several statistics that have a lower information overlap (on average) than other statistics, making these statistics prime candidates to capture unique information of a phylogenetic tree. Across empirical and simulated trees, we find that the eigenvector statistic and gamma statistic generally show very low information overlap with other statistics. While for empirical trees we find that statistics related to the Laplacian spectrum tend to contain a lot of unique information, for simulated trees, we find that the Pigot’s Rho, average ladder, max ladder and nLTT perform really well.

When restricted to a limited number of summary statistics, any of these statistics seem prime candidates to select as they show a restricted amount of overlap with other summary statistics.

An important aspect that is measured by many statistics, is the overall (im)balance of a phylogenetic tree. However, such statistics are typically defined and explored only for completely balanced or completely imbalanced trees and their relationship with changes in balance remains understudied. Here, we have introduced several algorithms to generate trees of intermediate balance in a systematic way. We found that most statistics do not vary linearly with balance, and for some statistics even found a non-monotonic relationship with balance, indicating that these statistics cannot be reliably used to compare balance of phylogenetic trees; these include the pitchforks, stairs, symmetry nodes, rogers, rquartet, equal weights Colless and average ladder. The non-linear relationship between balance and summary statistic value could also be due to the way we created trees of intermediate balance. Indeed, we found for some statistics strong variation across different imbalancing algorithms, e.g., for the maximum ladder statistic there is an almost linear relationship with balance if the intermediate balance is generated using terminal tips, starting with the youngest tips first, but a very non-linear relationship if random tips are picked. We conclude two things from our analysis of intermediate balance: firstly, almost all statistics have a non-linear or complicated relationship with balance, making interpretation of intermediate summary statistic values difficult. Secondly, the strongly differing relationships between summary statistic value and balance suggest that balance is not a one-dimensional property of a phylogenetic tree, but contains multidimensional information. This is in accordance with the previous subdivision into multiple correlated clusters of summary statistics we recovered for both empirical and simulated trees. The analysis of (im)balance of a phylogenetic tree therefore remains a complicated endeavor that is not easily defined except for at its extremes, and future explorations reaching into a better understanding of the relationship between balance statistics and intermediately balanced trees are highly warranted.

Across our analyses, we found that the different properties of the Laplacian spectrum often did not closely cluster together, but correlated with very different properties of phylogenetic trees, indicating that these measures of the Laplacian spectrum reflect different underlying properties: The principal eigenvalue typically clustered with balance statistics, whereas the asymmetry and peakedness clustered with branching time related statistics. The eigengap measure was a strong outlier for empirical trees, but clustered with branching time related statistics for simulated trees. Although the Laplacian spectrum related statistics are computationally slow and costly to compute in comparison with other statistics, the fact that these four properties of the spectrum correlate so differently makes the Laplacian spectrum related statistics interesting measures to be included in the phylogeneticists’ toolbox, although interpretation of its values may be difficult.

Previously, Tucker and colleagues identified three pillars of summary statistics (stemming from previous work in community assembly theory (Pavoine & Bonsall, 2011)): statistics measuring richness, divergence and regularity. They identified three ‘anchor’ statistics for these pillars: phylogenetic diversity, mean pair distance and variance in pair distance respectively. These three statistics rely heavily on including information from branch lengths, and in our results we recover that these statistics consistently cluster together amongst other branch length related statistics. This suggests that a large part of variation across trees is missed by selecting these ‘anchor’ statistics, and that inclusion of statistics that better represent topology would be warranted.

When using phylogenetic summary statistics to compare phylogenetic tree properties across different scenarios, models or taxonomic groups, selection of focal summary statistics is crucial to ensure a proper comparison. With our analysis here we have aimed to provide some guidance into which statistics to select. To reduce intercorrelation between statistics, we recommend to pick statistics from the different larger clusters. Furthermore, we recommend to pick at least one of the statistics that have been shown to show low correlation with all other statistics. When choosing statistics, picking multiple statistics out of those sets of statistics that are known to consistently correlate strongly with each other should be avoided. With these limitations in mind, it may be complicated to select the right combination of statistics, also taking into account that often intercorrelations of statistics may depend on the tree generating model, which may not always be known. Alternatively, we are confident that the computational speed of the treestats package can be leveraged by the researcher in order to explore a larger number of statistics and make an informed choice about summary statistic selection.

## Data availability

All scripts used for this manuscript are available on: https://github.com/thijsjanzen/treestats-scripts. The R package ‘treestats’ can be accessed on CRAN on : https://CRAN.R-project.org/package=treestats, and an ongoing development version can be found on: https://github.com/thijsjanzen/treestats.

## Author contributions

**Thijs Janzen**: Conceptualization, Methodology, Software, Formal Analysis, Writing – Original draft.

**Rampal S. Etienne:** Conceptualization, Methodology, Writing – Reviewing and editing

## Acknowledgements

We thank the Center for Information Technology of the University of Groningen for providing access to the Peregrine high-performance computing cluster.

## SUPPLEMENT

**Figure S1.**
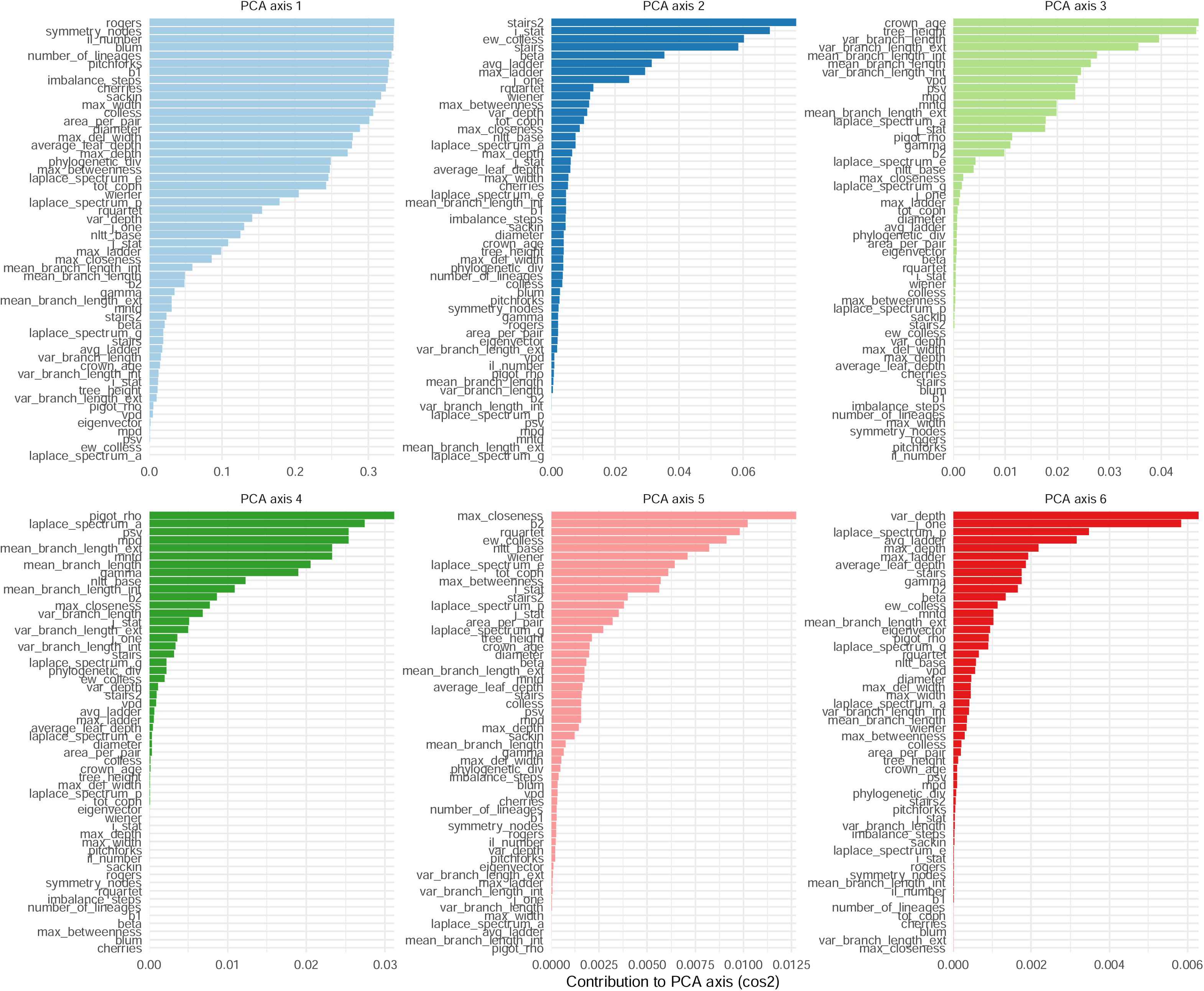
Contribution of each summary statistic to the first six PCA axes on 215 empirical phylogenetic trees.

### Time-variation of summary statistics

For every model, we considered 10,000 trees and for each we took the reconstructed tree up until time point *t*, and calculated all available summary statistics for this ‘subtree’. We did so for 100 timepoints equally spaced in [0, crown age]. We report the results (figure S2) relative to the crown age in order to facilitate comparison between trees of different crown age. Our results show clearly that the different diversification models show very different progressions through time, indicating that although some of these diversification models are nested, each model generates unique tree patterns. Furthermore, it shows, similar to Figure 4, that many tree statistics are unable to accurately correct for tree size, and that as a result, many patterns in trees are not constant throughout their age, possibly mainly due to their changing size (but see the cherries and pitchforks results).

**Figure S2.**
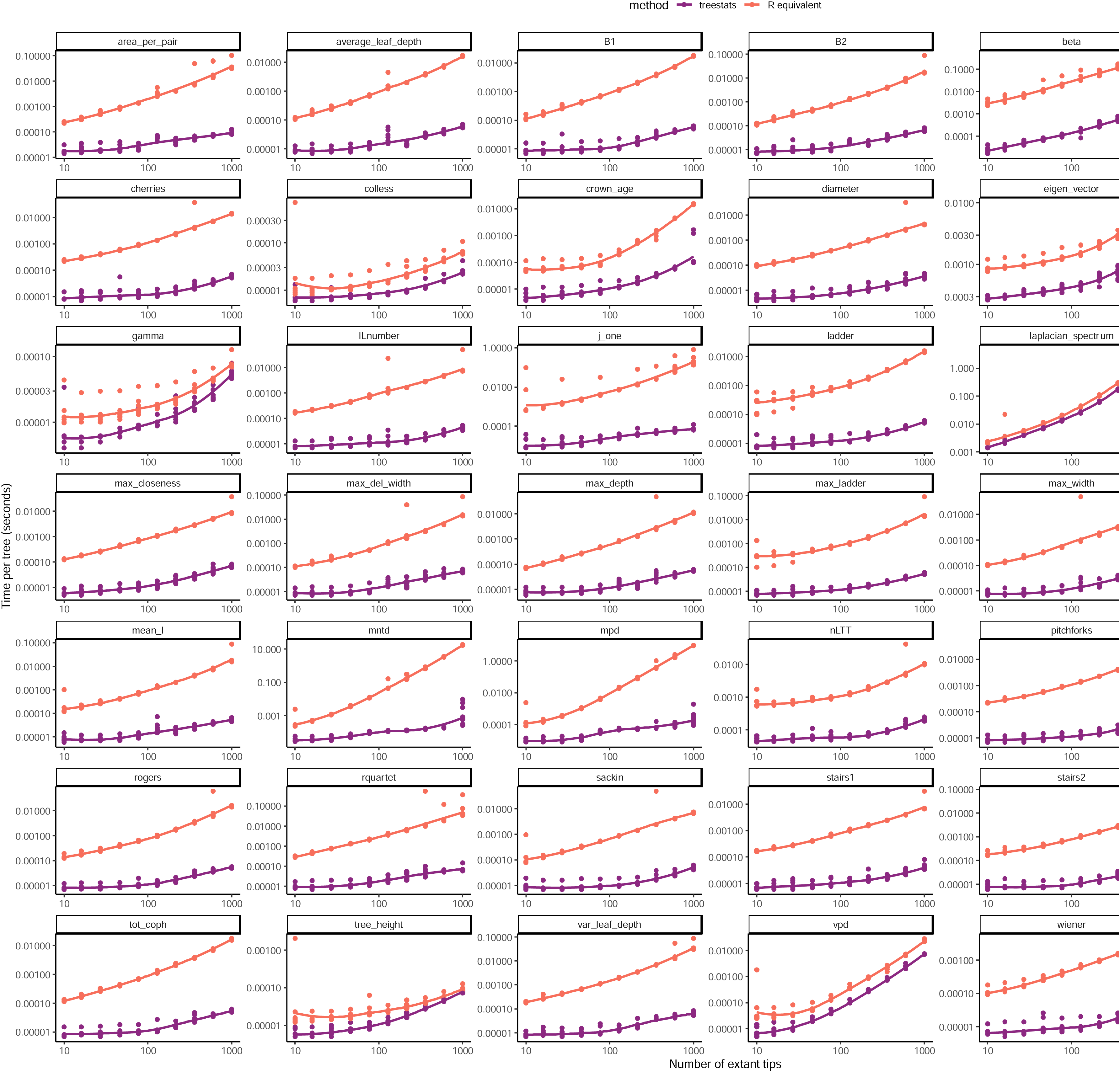
Comparison of computation speed (in seconds) of summary statistics included in the treestats package for which an equivalent R function is avaialbe. Comparison is with the fastest equivalent function in an available R package or R code. Shown are results for trees varying in the number of extant tips, per tree size, ten trees were generated using a Yule process with ape function rphylo. The line connecting the dots resembles the best fitting LOESS line as a helping guide. Performance measurements were completed on a MacBook Pro with M1 processor.

**Figure S3.**
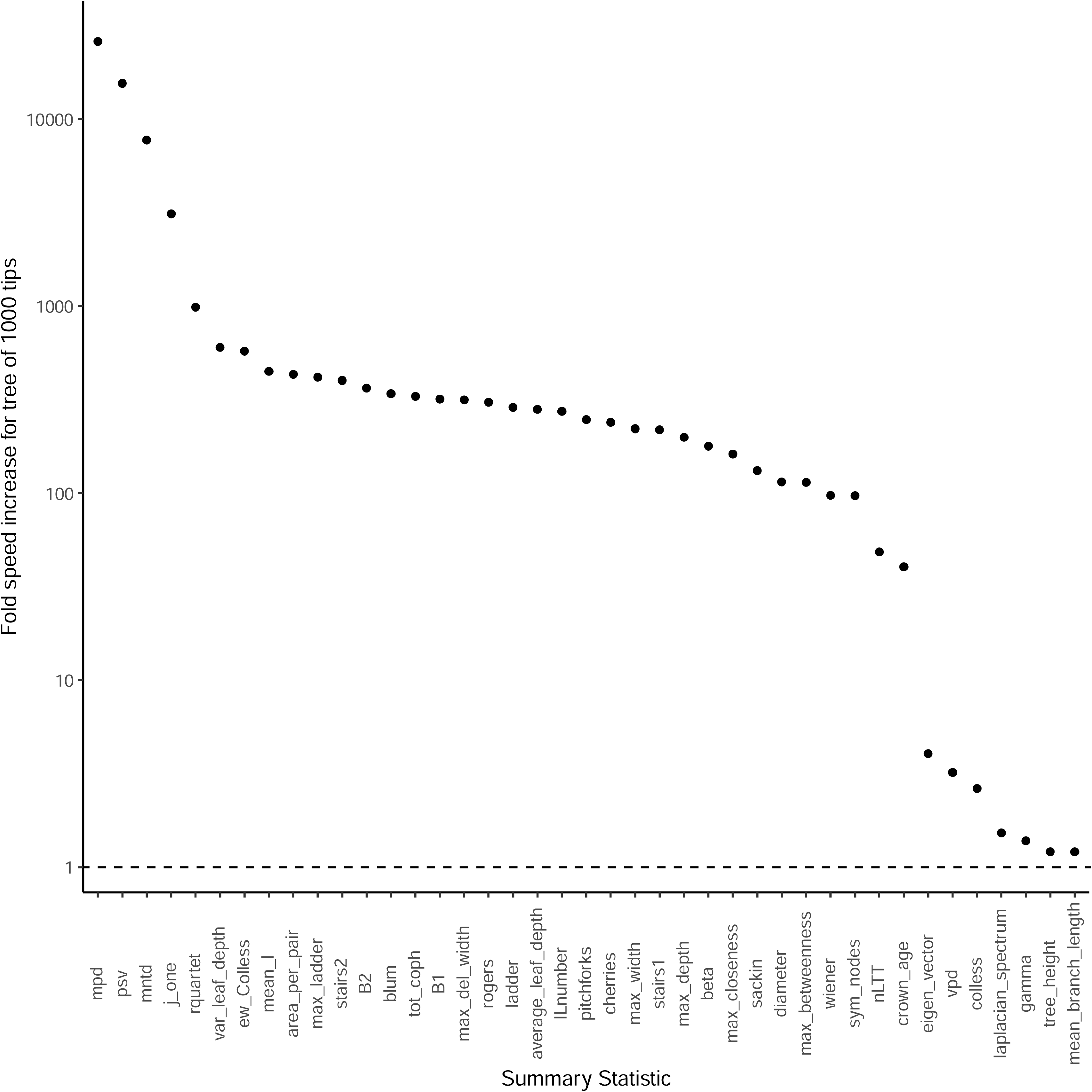
Relative speed increase (average old speed / average new speed) for all summary statistics, for a Yule tree with 1000 tips.

**Figure S4.**
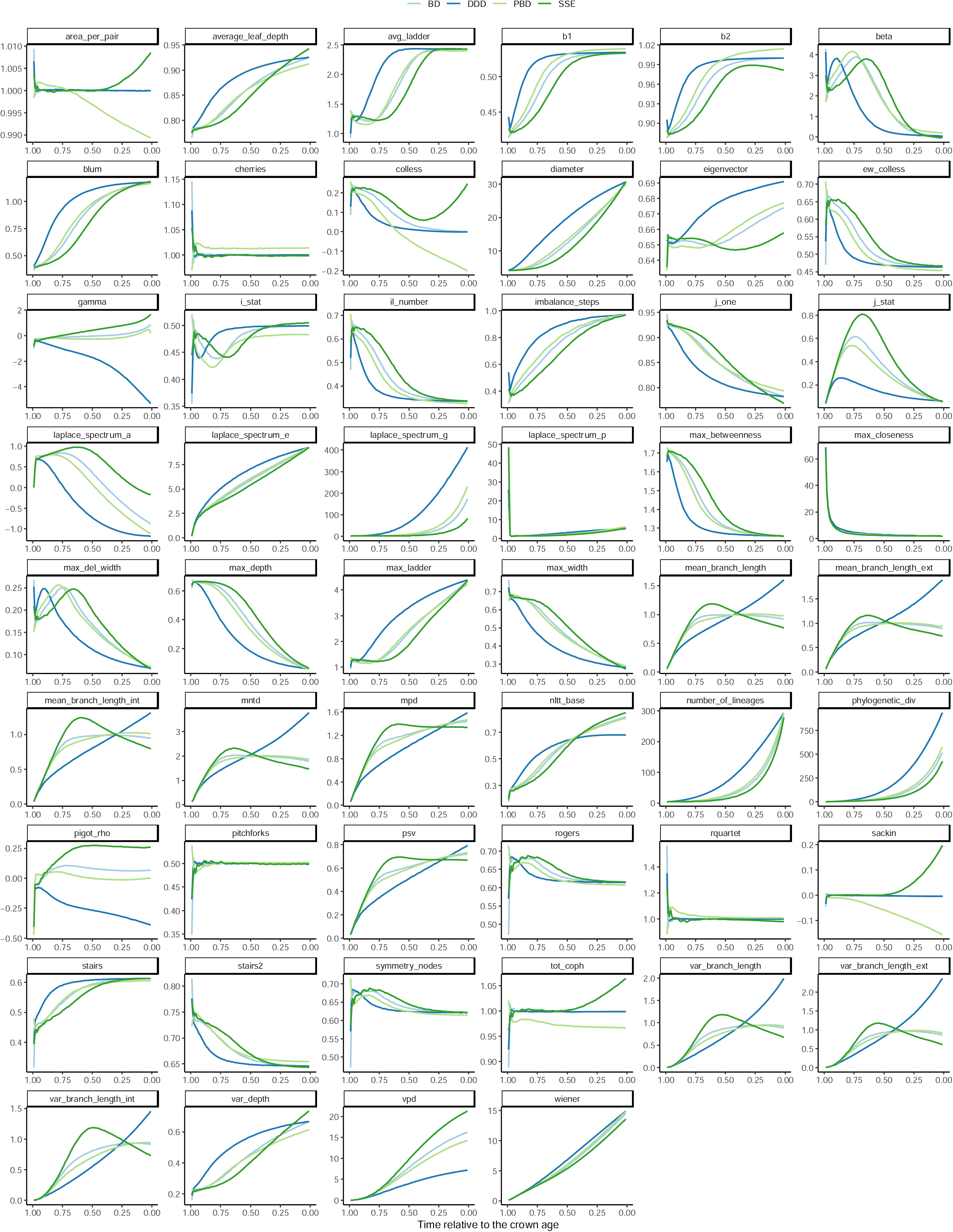
Summary statistics measured through time, relative to the crown age, with 1 indicating a value close to the crown, and 0 indicating a time close to the present, for the birth-death model, ddd model, pbd model and an sse model. Shown is the average across 10.000 trees per model.

## References

Aldous, D. (1996). Probability distributions on cladograms. Random Discrete Structures. http://link.springer.com/chapter/10.1007/978-1-4612-0719-1_1

Blum, M., & François, O. (2006). Which Random Processes Describe the Tree of Life? A Large-Scale Study of Phylogenetic Tree Imbalance. Systematic Biology, 55(4), 685–691. 10.1080/10635150600889625

Chindelevitch, L., Hayati, M., Poon, A. F. Y., & Colijn, C. (2021). Network science inspires novel tree shape statistics. PLOS ONE, 16(12), e0259877. 10.1371/journal.pone.0259877

Colijn, C., & Gardy, J. (2014). Phylogenetic tree shapes resolve disease transmission patterns. *Evolution*, Medicine and Public Health, 2014(1), 96–108. 10.1093/emph/eou018

Colless, D. (1982). Review of phylogenetics: The theory and practice of phylogenetic systematics. Systematic Zoology.

Condamine, F. L., Rolland, J., & Morlon, H. (2019). Assessing the causes of diversification slowdowns: Temperature-dependent and diversity-dependent models receive equivalent support. Ecology Letters, 22(11), 1900–1912. 10.1111/ele.13382

Coronado, T. M., Mir, A., Rosselló, F., & Rotger, L. (2020). On Sackin’s original proposal: The variance of the leaves’ depths as a phylogenetic balance index. BMC Bioinformatics, 21(1). 10.1186/s12859-020-3405-1

Coronado, T. M., Mir, A., Rosselló, F., & Valiente, G. (2019). A balance index for phylogenetic trees based on rooted quartets. Journal of Mathematical Biology, 79(3), 1105–1148. 10.1007/s00285-019-01377-w

Etienne, R. S., Haegeman, B., Stadler, T., Aze, T., Pearson, P. N., Purvis, A., & Phillimore, A. B. (2012). Diversity-dependence brings molecular phylogenies closer to agreement with the fossil record. Proceedings. Biological Sciences / The Royal Society, 279(1732), 1300–1309. 10.1098/rspb.2011.1439

Etienne, R. S., & Rosindell, J. (2012). Prolonging the past counteracts the pull of the present: Protracted speciation can explain observed slowdowns in diversification. Systematic Biology, 61(2), 204–213. 10.1093/sysbio/syr091

Faith, D. P. (1992). Conservation evaluation and phylogenetic diversity. Biological Conservation, 61(1), 1–10. 10.1016/0006-3207(92)91201-3

Fischer, M., Herbst, L., Kersting, S., Kuhn, L., & Wicke, K. (2021). Tree balance indices: A comprehensive survey (arXiv:2109.12281). arXiv.

FitzJohn, R. G. (2012). Diversitree: Comparative phylogenetic analyses of diversification in R. Methods in Ecology and Evolution, 3(6), 1084–1092. 10.1111/j.2041-210X.2012.00234.x

Fusco, G., & Cronk, Q. (1995). A new method for evaluating the shape of large phylogenies. Journal of Theoretical Biology, 235–243.

Guimarães Fabreti, L., & Höhna, S. (2023). Nucleotide Substitution Model Selection Is Not Necessary for Bayesian Inference of Phylogeny With Well-Behaved Priors. *Systematic Biology*, syad041. 10.1093/sysbio/syad041

Hagen, O., Hartmann, K., Steel, M., & Stadler, T. (2015). Age-Dependent Speciation Can Explain the Shape of Empirical Phylogenies. Systematic Biology, 64(3), 432–440. 10.1093/sysbio/syv001

Harmon, L. J., Schulte, J. A., Larson, A., & Losos, J. B. (2003). Tempo and mode of evolutionary radiation in iguanian lizards. Science, 301(5635), 961–964. 10.1126/science.1084786

Helmus, M. R., Savage, K., Diebel, M. W., Maxted, J. T., & Ives, A. R. (2007). Separating the determinants of phylogenetic community structure. Ecology Letters, 10(10), 917–925. 10.1111/j.1461-0248.2007.01083.x

Herrera-Alsina, L., van Els, P., & Etienne, R. S. (2018). Detecting the Dependence of Diversification on Multiple Traits from Phylogenetic Trees and Trait Data. Systematic Biology, 0(0), 1–12. 10.1093/sysbio/syy057

Izsák, J., & Papp, L. (2000). A link between ecological diversity indices and measures of biodiversity. Ecological Modelling, 130(1), 151–156. 10.1016/S0304-3800(00)00203-9

Janzen, T., & Etienne, R. S. (2017). Inferring the role of habitat dynamics in driving diversification: Evidence for a species pump in Lake Tanganyika cichlids. bioRxiv, 085431. 10.1111/cobi.12609.This

Janzen, T., Höhna, S., & Etienne, R. S. (2015). Approximate Bayesian Computation of diversification rates from molecular phylogenies: Introducing a new efficient summary statistic, the nLTT. Methods in Ecology and Evolution, 6, 566–575. 10.1111/2041-210X.12350

Kendall, M., Boyd, M., & Colijn, C. (2018). phyloTop: Calculating Topological Properties of Phylogenies [Computer software].

Kersting, S. J., & Fischer, M. (2021). Measuring tree balance using symmetry nodes—A new balance index and its extremal properties. Mathematical Biosciences, 341. 10.1016/j.mbs.2021.108690

Lambert, S., Voznica, J., & Morlon, H. (2023). Deep Learning from Phylogenies for Diversification Analyses. *Systematic Biology*, syad044. 10.1093/sysbio/syad044

Lemant, J., Le Sueur, C., Manojlović, V., & Noble, R. (2022). Robust, Universal Tree Balance Indices. Systematic Biology, 71(5), 1210–1224. 10.1093/sysbio/syac027

Lewitus, E., & Morlon, H. (2016). Characterizing and comparing phylogenies from their laplacian spectrum. Systematic Biology, 65(3), 495–507. 10.1093/sysbio/syv116

Lima, T. A., Marquitti, F. M. D., & de Aguiar, M. A. M. (2020). Measuring Tree Balance with Normalized Tree Area. http://arxiv.org/abs/2008.12867

Liow, L. H., Quental, T. B., & Marshall, C. R. (2010). When Can Decreasing Diversification Rates Be Detected with Molecular Phylogenies and the Fossil Record? Systematic Biology, 59(6), 646–659. 10.1093/sysbio/syq052

McKenzie, A., Steel, M., & McKenzie, A. (1999). Distributions of cherries for two models of trees.

Mir, A., Rosselló, F., & Rotger, L. a. (2013). A new balance index for phylogenetic trees. Mathematical Biosciences, 241(1), 125–136. 10.1016/j.mbs.2012.10.005

Mooers, A. O. & S. B. Heard. (1997). Macroevolution and the shape of phylogenetic trees. The Quarterly Review of Biology, 72, 31–54.

Nee, S. (2001). Inferring speciation rates from phylogenies. Evolution, 55(4), 661–668.

Nee, S., & Holmes, E. (1994). Extinction rates can be estimated from molecular phylogenies. Philosophical Transactions: Biological Sciences, 344(1307), 77–82. 10.1111/j.1558-5646.2009.00926.x

Norström, M. M., Prosperi, M. C. F., Gray, R. R., Karlsson, A. C., & Salemi, M. (2012). PhyloTempo: A set of R scripts for assessing and visualizing temporal clustering in genealogies inferred from serially sampled viral sequences. Evolutionary Bioinformatics, 2012(8), 261–269. 10.4137/EBO.S9738

O’Meara, B., Sanchez-Reyes, L. L., Eastman, J., Heath, T., Wright, A., Schliep, K., Chamberlain, S., Midford, P., Harmon, L., Brown, J., Pennell, M., Alfaro, M., & McTavish, E. J. (2023). datelife: Scientific Data on Time of Lineage Divergence for Your Taxa. 10.5281/zenodo.593938

Pagel, M., Meade, A., & Barker, D. (2004). Bayesian Estimation of Ancestral Character States on Phylogenies. Systematic Biology, 53(5), 673–684. 10.1080/10635150490522232

Pavoine, S., & Bonsall, M. B. (2011). Measuring biodiversity to explain community assembly: A unified approach. Biological Reviews, 86(4), 792–812. 10.1111/j.1469-185X.2010.00171.x

Pigot, A. L., Phillimore, A. B., Owens, I. P. F., & Orme, C. D. L. (2010). The shape and temporal dynamics of phylogenetic trees arising from geographic speciation. Systematic Biology, 59(6), 660–673. 10.1093/sysbio/syq058

Purvis, A., Katzourakis, A., & Agapow, P.-M. (2002). Evaluating phylogenetic tree shape: Two modifications to Fusco & Cronk’s method. Journal of Theoretical Biology, 214(1), 99–103. 10.1006/jtbi.2001.2443

Pybus, O., & Harvey, P. (2000). Testing macro–evolutionary models using incomplete molecular phylogenies. Proceedings of the Royal Society B: Biological Sciences, 267(1459), 2267–2272. 10.1098/rspb.2000.1278

Revell, L. J. (2012). phytools: An R package for phylogenetic comparative biology (and other things). Methods in Ecology and Evolution, 3, 217–223. 10.1111/j.2041-210X.2011.00169.x

Revell, L. J., & Harmon, L. J. (2022). Phylogenetic Comparative Methods in R. Princeton University Press.

Richter, F., Janzen, T., Hildenbrandt, H., Wit, E. C., & Etienne, R. S. (2021). Detecting phylodiversity-dependent diversification with a general phylogenetic inference framework (p. 2021.07.01.450729). bioRxiv. 10.1101/2021.07.01.450729

Rogers, J. S. (1996). CENTRAL MOMENTS AND PROBABILITY DISTRIBUTIONS OF THREE MEASURES OF PHYLOGENETIC TREE IMBALANCE. In Losos and Adler (Vol. 45, pp. 99–110). Rogers. https://academic.oup.com/sysbio/article/45/1/99/1631748

Ruffley, M., Peterson, K., Week, B., Tank, D. C., & Harmon, L. J. (2019). Identifying models of trait-mediated community assembly using random forests and approximate Bayesian computation. Ecology and Evolution, 9(23), 13218–13230. 10.1002/ece3.5773

Sackin, M. (1972). “Good” and “bad” phenograms. Systematic Biology, 21(2), 225–226.

Saulnier, E., Gascuel, O., & Alizon, S. (2017). Inferring epidemiological parameters from phylogenies using regression-ABC: A comparative study. PLOS Computational Biology, 13(3), e1005416. 10.1371/journal.pcbi.1005416

Shao, K.-T., & Sokal, R. R. (1990). Tree Balance. Systematic Biology, 39(3), 266–276. 10.2307/2992186

Tsirogiannis, C., Sandel, B., & Cheliotis, D. (2012). Efficient Computation of Popular Phylogenetic Tree Measures. In B. Raphael & J. Tang (Eds.), Algorithms in Bioinformatics (Vol. 7534, pp. 30–43). Springer Berlin Heidelberg. 10.1007/978-3-642-33122-0_3

Tucker, C. M., Cadotte, M. W., Carvalho, S. B., Jonathan Davies, T., Ferrier, S., Fritz, S. A., Grenyer, R., Helmus, M. R., Jin, L. S., Mooers, A. O., Pavoine, S., Purschke, O., Redding, D. W., Rosauer, D. F., Winter, M., & Mazel, F. (2017). A guide to phylogenetic metrics for conservation, community ecology and macroecology. Biological Reviews, 92(2), 698–715. 10.1111/brv.12252

Verboom, G. A., Boucher, F. C., Ackerly, D. D., Wootton, L. M., & Freyman, W. A. (2020). Species Selection Regime and Phylogenetic Tree Shape. Systematic Biology, 69(4), 774–794. 10.1093/sysbio/syz076

Voznica, J., Zhukova, A., Boskova, V., Saulnier, E., Lemoine, F., Moslonka-Lefebvre, M., & Gascuel, O. (2022). Deep learning from phylogenies to uncover the epidemiological dynamics of outbreaks. Nature Communications, 13(1), Article 1. 10.1038/s41467-022-31511-0

Webb, C. O., Ackerly, D. D., McPeek, M. A., & Donoghue, M. J. (2002). Phylogenies and community ecology. Annual Review of Ecology and Systematics, 33, 475–505. 10.1146/annurev.ecolsys.33.010802.150448

Yu, Y., Harris, A. J., Blair, C., & He, X. (2015). RASP (Reconstruct Ancestral State in Phylogenies): A tool for historical biogeography. Molecular Phylogenetics and Evolution, 87, 46–49. 10.1016/j.ympev.2015.03.008

